# Foliar fungi alter reproductive timing and allocation in *Arabidopsis* under normal and water-stressed conditions

**DOI:** 10.1101/519678

**Authors:** Geoffrey Zahn, Anthony S. Amend

## Abstract

Microbes influence plant phenotypes but most known examples of this are from the study of below-ground microbes and plant disease modification. To examine the potential importance of phyllosphere microbes on non-disease related plant traits, we used sterile *Arabidopsis* clones to test the effects of foliar fungi on flowering phenology and reproductive allocation under conditions of varying water stress. We inoculated the sterile plants with fully-factorial combinations of four fungal isolates, then measured flowering time and reproductive allocation for each treatment group under normal and water-stressed conditions. All plants inoculated with foliar fungi had significantly later flowering and greater seed mass than the sterile control groups. The magnitude of this effect depended on the specific fungi present, but individual fungal effects diminished as inoculum richness increased. Above-ground microbes likely influence other plant traits as well and should be considered in any study measuring plant phenotypes.

## INTRODUCTION

Plants are hosts to a diverse array of endophytic and epiphytic microorganisms that, together with the plant, make up the holobiont. Under natural conditions, many “plant” traits depend entirely, or in part, on members of the plant microbiome (Vandenkoornhuyse *et al.*, 2015). However, the more we discover about the holobiont, the more questions arise about the relative roles of microbial community dynamics, plant genotype, and environmental factors that shape a plant’s extended phenotype (Preston, 2017).

Untangling the processes of plant-microbe interactions can be very complicated, with plants shaping their microbiome and the microbes, in turn, shaping attributes of plant phenotype (Mendes *et al.*, 2014; Edwards *et al.*, 2015). Work that has simplified these systems by incorporating sterile model plants and manipulating the microbial constituents is, however, beginning to reveal a much needed mechanistic understanding of plant microbiome assembly and function (Panke-Buisse *et al.*, 2015; Wolfe, 2018).

The influence of microbes on plant traits can be subtle or dramatic. For example, that bacterial root endophytes are known to modify plant phenotypes in some cases by changing when and how carbon is allocated within plant tissues (Henning *et al.*, 2016), and that transplantations of natural soil biota to gnotobiotic plants can alter flowering phenology (Wagner *et al.*, 2014). Although most work has focused on below-ground microbes and plant tissues (Rossmann *et al.*, 2017), it is clear that microbes associated with above-ground (phyllosphere) tissues also shape plant traits.

Aside from modifying plant disease (Falconi & Mendgen, 1994; Arnold *et al.*, 2003; Busby *et al.*, 2015), phyllosphere-associated microbes alter plant traits such as leaf wettability and xylem conductivity, (Beattie, 2011), cuticle permeability and transpiration (Ritpitakphong *et al.*, 2016), and even the biosynthesis of plant hormones (Egamberdieva *et al.*, 2017). Additionally, above-ground microbes can influence seed mass (Saari *et al.*, 2010), leaf size (Davitt *et al.*, 2000), frost sensitivity (Bertrand *et al.*, 2007), and shoot height (Perrine-Walker *et al.*, 2007). No doubt, many more microbially-mediated plant traits await discovery. An increased understanding of phyllosphere microbial communities and which plant traits they can alter is crucial for informing emergent agricultural (Schlaeppi & Bulgarelli, 2014), industrial (Doty, 2017), and conservation (Zahn & Amend, 2017) practices.

In this study we investigated how foliar fungal endophytes impact flowering phenology and seed mass, important ecological traits (Westoby *et al.*, 1992; Rosas *et al.*, 2014) under normal and water-stressed conditions. Using gnotobiotic clones of *Arabidopsis thaliana* inoculated with a fully-factorial combination of four fungal isolates with varying phylogenetic distance from each other, we sought to uncover whether the presence and identity of foliar fungi affect reproductive timing and allocation under normal and stressed conditions.

## MATERIALS AND METHODS

### Overview

Replicates of sterile *Arabidopsis* seedlings were inoculated with factorial combinations of 4 fungal isolates having varying phylogenetic distance from each other. The experiment was duplicated under normal (surface soil never allowed to completely dry) and water-stressed (soil surface allowed to dry for 2 days between waterings) conditions. Flowering phenology (days to first flower) and average seed mass were measured.

### Plant selection and growth

Arabidopsis bulk germplasm was generated from a single line obtained from TAIR (www.arabidopsis.org, Accession: CS22468). Seeds were surface sterilized by shaking in 10% bleach containing 0.05% Tween 20 detergent for 10 minutes, 70% ethanol for 2 minutes, 3 rinses of sterile DI water. After 2 weeks cold treatment at 4deg C, they were sown in autoclaved soil-less potting medium (Sunshine #4, Sun Grow, Chicago, IL). Seeds were germinated in the dark at 21deg C for one week, then transplanted into 3” pots of sterile potting medium (Day 0) and grown in a Percival growth chamber at 20 deg C with 16h/8h day/night cycle. Successfully-germinated seedlings were randomly assigned to fungal and water treatments, so that sterile controls and each factorial treatment combination had 5 replicate individuals. Replicates from each group were kept together in the growth chamber to minimize the potential for cross-contamination and group locations were rotated each day. Plants were watered from below with sterile DI water when the soil surface began to dry (for normal water treatment) and 2 days after the soil surface dried (water stress treatment). The first and third waterings contained 15-5-15 fertilizer at [200 ppm N]. Plants were not allowed to sit in standing water. The date of first flowering was recorded for each plant and waterings were halted 7 days after 80% of plants in each group initiated flowering. At the end of flowering, seeds were collected from each individual and dried at 40deg C for two weeks, after which the total mass for 50 randomly selected seeds from each individual was measured to determine the mean mass of a single seed from each plant.

### Fungal isolate inoculum preparation and inoculation

Fungi were originally isolated from Brassicaseae (kale and broccoli cultivars grown in an outdoor garden) leaf surfaces by running a moistened sterile swab across leaf surfaces and then streaking it onto MEA agar (Malt extract [1 g/L], Yeast extract [1g/L], Agar [10g/L]) amended with Streptomycin [20mg/mL] and Kanamycin [20mg/mL]. Morphologically distinct fungal colonies were serially axenized on MEA plates. DNA was extracted from 8 fungal isolates using the Extract-n-Amp protocol (Sigma-Aldrich) and subjected to bi-directional Sanger sequencing of the ITS1-28S region using the primers ITS1F (5’-CTTGGTCATTTAGAGGAAGTAA-3’) and TW-13 (5’-GGTCCGTGTTTCAAGACG-3’). Taxonomy was assigned by comparing consensus sequences against the NCBI nucleotide database using BLAST. Phylogenetic relatedness was estimated as the sum of patristic branch lengths from a Jukes-Cantor model neighbor-joining tree of the 5.8S rDNA region containing 70 other fungi (Cullings & Vogler, 1998).

Four sporulating fungal isolates with varying levels of phylogenetic distance (SI Table 1) were chosen as inoculum sources. Phylogenetic placement was consistent with the top BLAST hit. We refer to the fungal taxa as A-D within this paper but taxonomic assignments are reported in Table 1. Spores from isolates A, B, C, and D were scraped from the surfaces of cultures and centrifugally washed three times in sterile water (500 RCF for 10 minutes, supernatant removed from pellet, pellet resuspended in 1mL sterile water). Spore concentrations were quantified and normalized using a hemocytometer and DIC microscopy at 40X magnification before diluting them into 200 mL of 0.1% sterile agarose solution. Each fungal inoculum had an equivalent final spore concentration of 6,944 cells/mL, regardless of the number of species present. Negative controls consisted of a heat-killed combination of all 4 fungal isolates at 6,944 cells/mL in sterile 0.1% agarose solution.

**Table 1.** Taxonomic assignments for the four fungal isolates used in this study. Sequences have been deposited at NCBI under the GenBank Accession Numbers given. Taxonomy was chosen based on the top named BLAST hit against the NCBI nr database.

| <u>Isolate ID</u> | <u>GenBank</u><br><u>Accession</u><br><u>No.</u> | <u>Top BLAST hit</u> | <u>Tentative Taxonomic Assignment</u> |
| --- | --- | --- | --- |
| **A** | **<u>MH972194</u>** | NR_157531.1 | *Pleospora rosae* |
| **B** | **<u>MH972195</u>** | KX641964.1 | *Cochliobolus sp.* |
| **C** | **<u>MH972196</u>** | KX664408.1 | *Alternaria tenuissima* |
| **D** | **<u>MH972197</u>** | KX664406.1 | *Cladosporium macrocarpum* |

After the first true leaves appeared on at least 75% of all germinated plants (Day 6) fungal inoculae were sprayed to fully cover plants and soil surfaces. Plants were sprayed weekly with their respective fungal treatments until rosettes reached full size (Day 23).

### Fungal isolate growth rates

Three replicate cultures were prepared for isolates in which fungi were grown singly and in each possible combination with each other. Agar blocks (0.25 cm2) of vegetative mycelium was cut from pure cultures and placed onto the surface of MEA agar and grown at ambient temperature for 9 days. To measure baseline and competitive growth rates, replicates of pure cultures and each factorial combination were photographed and mycelium surface area was measured with ImageJ (Supporting Info) on days 4 and 9. Growth rates were determined as the increase in mycelial surface area over 5 days. Growth rates of isolates in competition were relativized against pure culture growth rates.

### Statistics

All analyses were performed in R (version 3.3.3). An analysis of variance (ANOVA) test was used to determine the significance of fungal treatment and water stress on flowering time. Flowering time was regressed onto seed mass (a proxy for plant fitness) with a linear model to determine the relationship between phenology and fitness. To estimate the role of each of the four fungal isolates in altering flowering time, we used a general linear model with community matrix components as predictors. The intercept was excluded and Type-III Sums of Squares were obtained with the *car* package (Fox & Weisberg, 2011). Figures were constructed with the *ggplot2* package (Wickham, 2009).

## RESULTS

### Flowering phenology and seed mass

All individuals inoculated with viable fungal isolates flowered later and had larger seeds than sterile controls, irrespective of watering levels (Fig. 1; SI Table 2). We observed significant changes to flowering phenology depending on fungal treatment group (ANOVA: F_128, 15_= 36.734; P < 2^-16^) and water regime (ANOVA: F_128, 1_ = 162.438; P < 2^-16^).

**Fig. 1.**
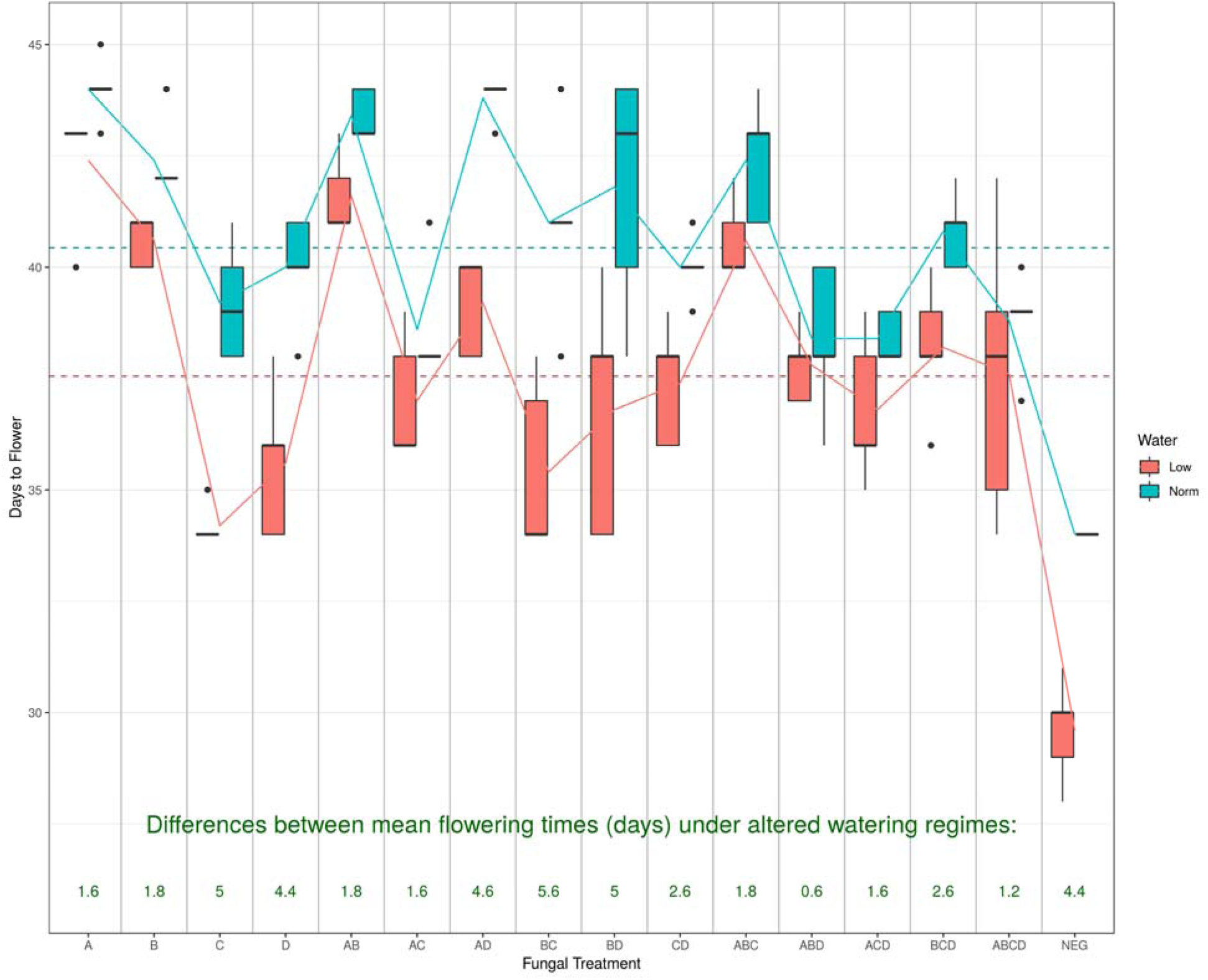
Flowering time (days to first flower emergence) for each treatment. Water-stressed plants (shown in red) had consistently shorter flowering times than plants under normal water conditions (shown in blue). The mean number of days in flowering time reduction (shown in green) refers to the difference between normal and water-stressed flowering times for each group. Fungal treatment groups refer to all combinations of the 4 fungal isolates.

Water stress consistently led to earlier flowering in all treatment groups, including controls. The magnitude of this change, however, varied between fungal community treatments (ANOVA: F_128, 15_ = 3.292; P = 0.000118; SI Table 2). The average reduction in flowering time due to water stress was 1.6 days for plants treated with isolate A, 1.8 days for those treated with isolate B, 5 days for plants treated with isolate C, and 4.4 days for those treated with isolate D. A related pattern was observed for these four isolates in their effects on plants’ overall watering times. Plants inoculated with isolate A flowered later on average than plants inoculated with isolates C or D, regardless of water stress (SI Table 3).

Inoculum community richness had no influence on the observed lengthening of flowering time (SI Fig. 1). Even the presence of a single fungal isolate, regardless of identity, increased the mean flowering time over sterile control plants (Fig. 2).

**Fig. 2.**
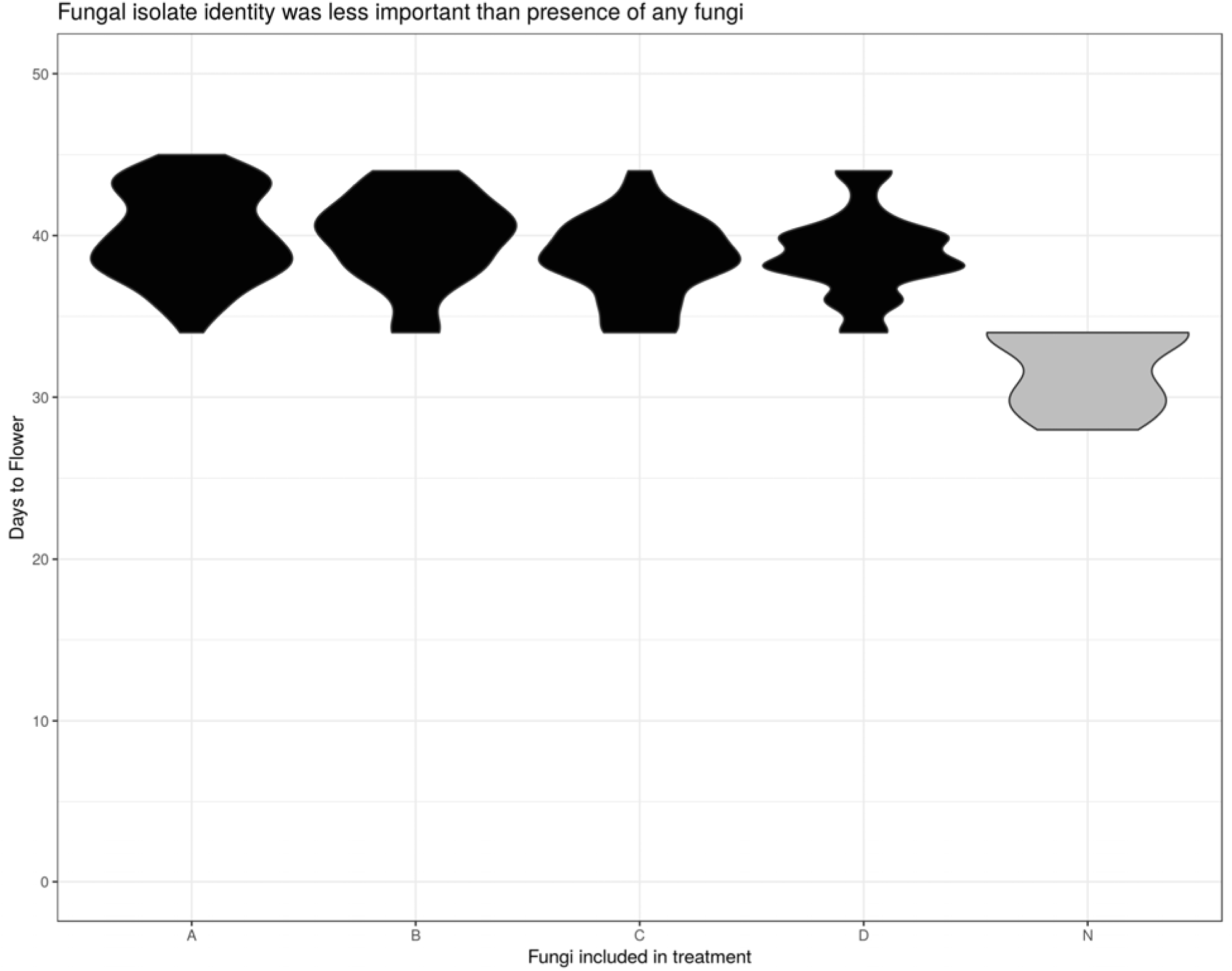
The presence of any specific fungal isolate was less importance than the presence of any fungus, regardless of identity. X-axis shows the presence of a given fungal isolate in a treatment group (N refers to sterile controls with no fungi). Y-axis shows violin distributions of flowering times (in days) for plant treated with that fungus in any combination of other inoculae.

Mean seed mass was positively correlated with later flowering times (F_1, 123_ = 22.24; P < 0.0005; Adj. R-sq = 0.1462; SI Fig. 2).

### Fungal isolate growth rates

The culture-based experiment allowed us to see how the four fungal isolates behaved in combination with each other (Fig. 3). Isolate B was a poor competitor under these conditions, often growing at half (or less) of its baseline rate when in competition with any or all of the other isolates. In culture, it seemed to facilitate faster growth in the other three isolates. The patterns we found in culture, however, had little predictive power for results on plants.

**Fig. 3.**
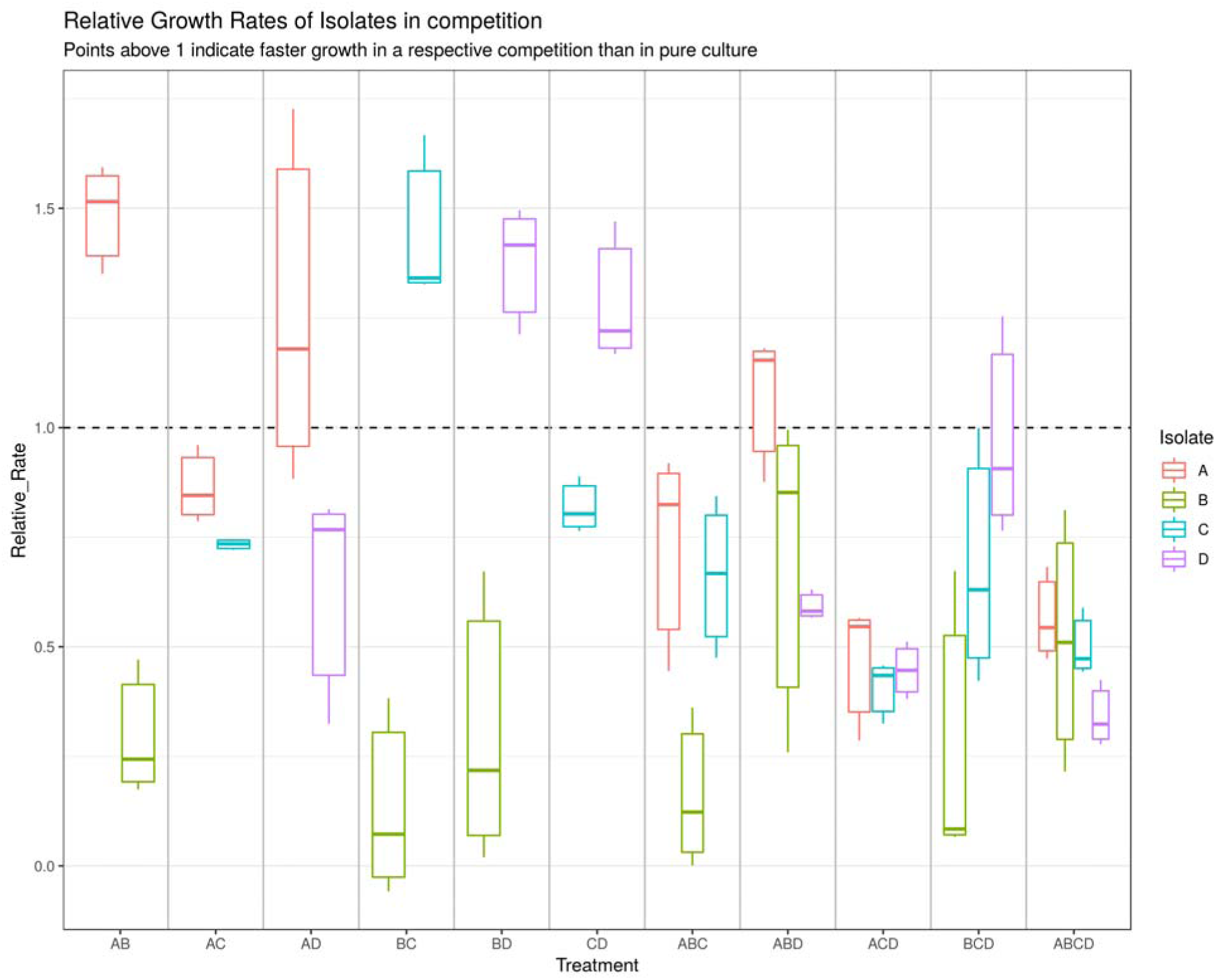
Fungal competitive growth rates on MEA medium as the proportion of growth rates in pure culture (cm2 day-1). X-axis = competition groups; Y-axis = Proportion of pure culture growth rate. Colored by fungal isolate. Values above 1.0 denote fungi that outperformed baseline growth rates in the presence of other isolate(s).

## DISCUSSION

Flowering timing is strongly determined by genetic (Rosas *et al.*, 2014), and environmental (Banta *et al.*, 2012) factors but, like other plant traits, it is also dependent on microbial members of the holobiont. Prior efforts have demonstrated important roles for soil bacteria in flowering phenology and drought response (Lau & Lennon, 2012; Wagner *et al.*, 2014). Here, we show that above-ground fungal members of the holobiont can also modify flowering phenology. Interestingly, at least for the isolates tested in this study, fungal identity was much less important than the fact that fungi were present.

In general, plants receiving isolates A and/or B (putatively assigned to *Pleospora rosae* and *P. herbarum*, respectively) had later flowering times and a reduction in the deleterious effects of drought compared with plants receiving isolates C and/or D (putatively assigned to *Alternaria tenuissima* and *Cladosporium macrocarpum*, respectively). However, in the higher-order interactions when more than two fungal strains were present in an inoculum, these patterns were not always maintained.

There was also an interaction between water stress and the functional benefits of harboring foliar fungi. Resources available to the phyllosphere microbial community may alter or even erase the advantages of that community for a plant (Berg & Koskella, 2018). Here we observe that drought conditions consistently resulted in earlier flowering times and reduced seed mass, but the magnitude of this effect depended on the fungal community present. Again, no consistent pattern could be found with respect to any specific isolate due to complexities in higher-order interactions.

Culture-based competition results shed little light on why simple interactions were not predictive of higher-order results. For example, both isolates A and B were associated with later flowering time individually and together, and isolate D was associated with earlier flowering time, especially under water stress. However, a mixed inoculum of A, B, and D showed no significant changes in flowering time from the mean of all treatment groups. In culture, the growth rate of isolate D was reduced by both isolates A and B, and the growth rate of isolate A was increased in the presence of isolate D (Fig. 3). This led us to predict that, in the presence of isolates A and B, the growth of isolate D would be reduced and it’s solitary effect on flowering time would be less pronounced, but this was not seen. This shows that there isn’t a clear relationship between competitiveness in a rich culture medium and effects on plant phenotype. Something more complex is taking place on the surface of plant leaves.

This higher-order unpredictability is not surprising, given that fungal community interactions and assembly processes are still not well understood. Our system was very simple, with just four fungal isolates. In natural systems, we must consider that determinants of microbial colonization of a given plant tissue include host plant identity and attributes, local abiotic conditions, and microbe-microbe interactions (Aleklett *et al.*, 2014). Of further importance are: priority effects (Tucker & Fukami, 2014; Toju *et al.*, 2018), phylogenetic relatedness and trait conservatism (Maherali & Klironomos, 2007), environmental filtering (Glassman *et al.*, 2017; Whitman *et al.*, 2018), and the interactions between these processes (Peay *et al.*, 2011). The specific mechanism by which fungi postponed flowering time in this study is also not known. Foliar fungi may alter photosynthetic efficiency, transpiration, and water use efficiency, particularly when plants are experiencing stress (Pinto *et al.*, 2000; Li *et al.*, 2012). They also produce abundant secondary metabolites that have received little attention outside of pathology applications (Suryanarayanan, 2013), which may be important for flowering phenology.

While we currently lack a detailed mechanistic explanation for why some fungal communities generated a greater effect than others, all the foliar fungal communities tested resulted in plants with significantly later flowering times and greater seed mass than the sterile controls. This, along with other research demonstrating that belowground microbes have strong influences on plant phenotype, should add complexity to the interpretation of “plant” traits, which might actually be microbial traits in disguise.

## Supporting information

Raw Data and analysis code

## ACKNOWLEDGEMENTS

The authors thank Benjamin Hoyt for assistance with experimental monitoring. This project was funded through the US Army cooperative agreement W9126G-11-2-0066 with Pacific Cooperative Studies Unit and the NSF DEB-1255972.

## AUTHOR CONTRIBUTIONS

AA and GZ conceived and designed the experiments. AA provided materials, reagents, and equipment. GZ performed the experiments. AA and GZ analyzed data and wrote the manuscript.

## DATA ACCESSIBILITY STATEMENT

All raw data are provided along with analysis code in the Supporting Information. DNA sequences for fungal isolates used in this study have been uploaded to NCBI and are given in Table 1.

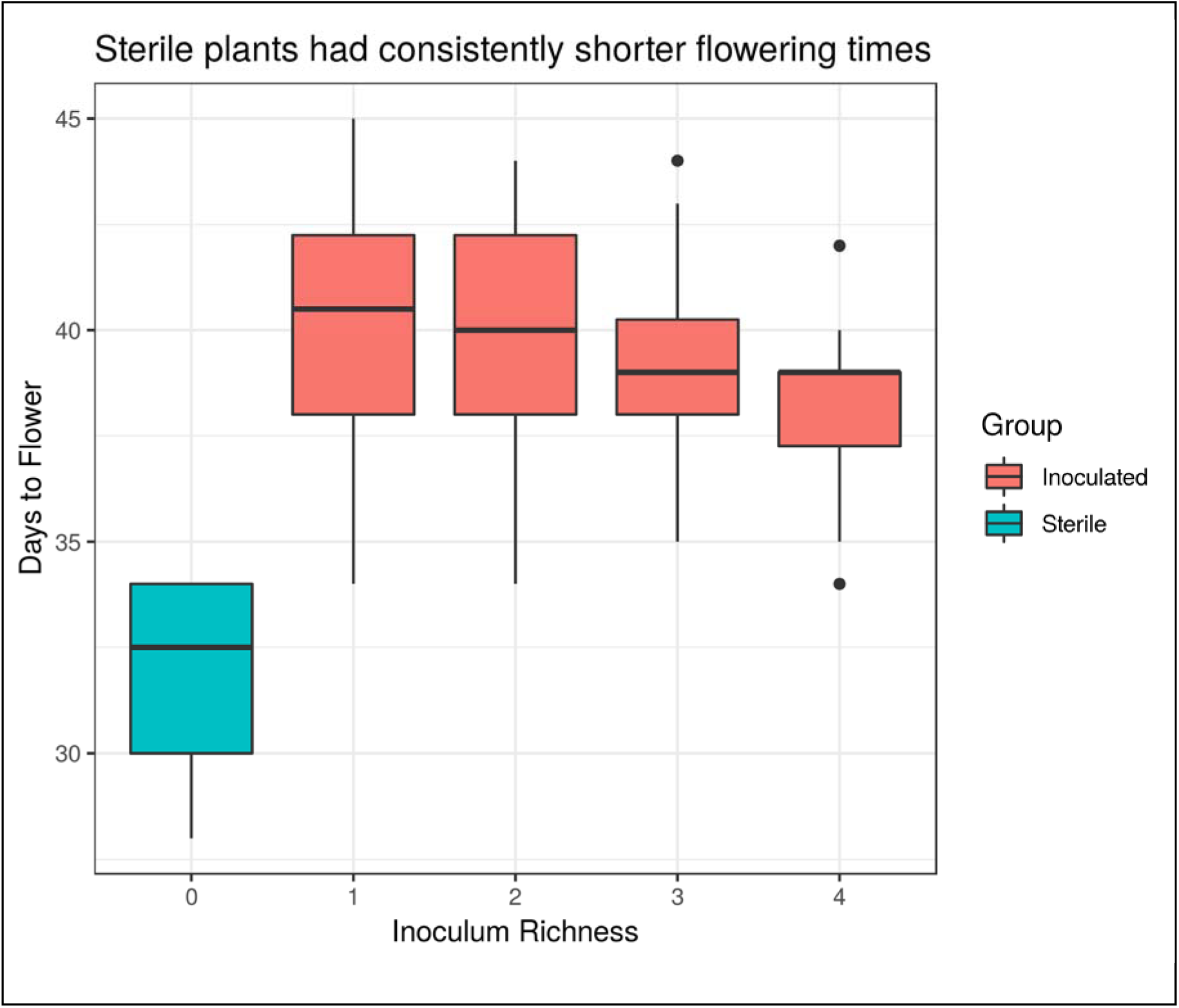

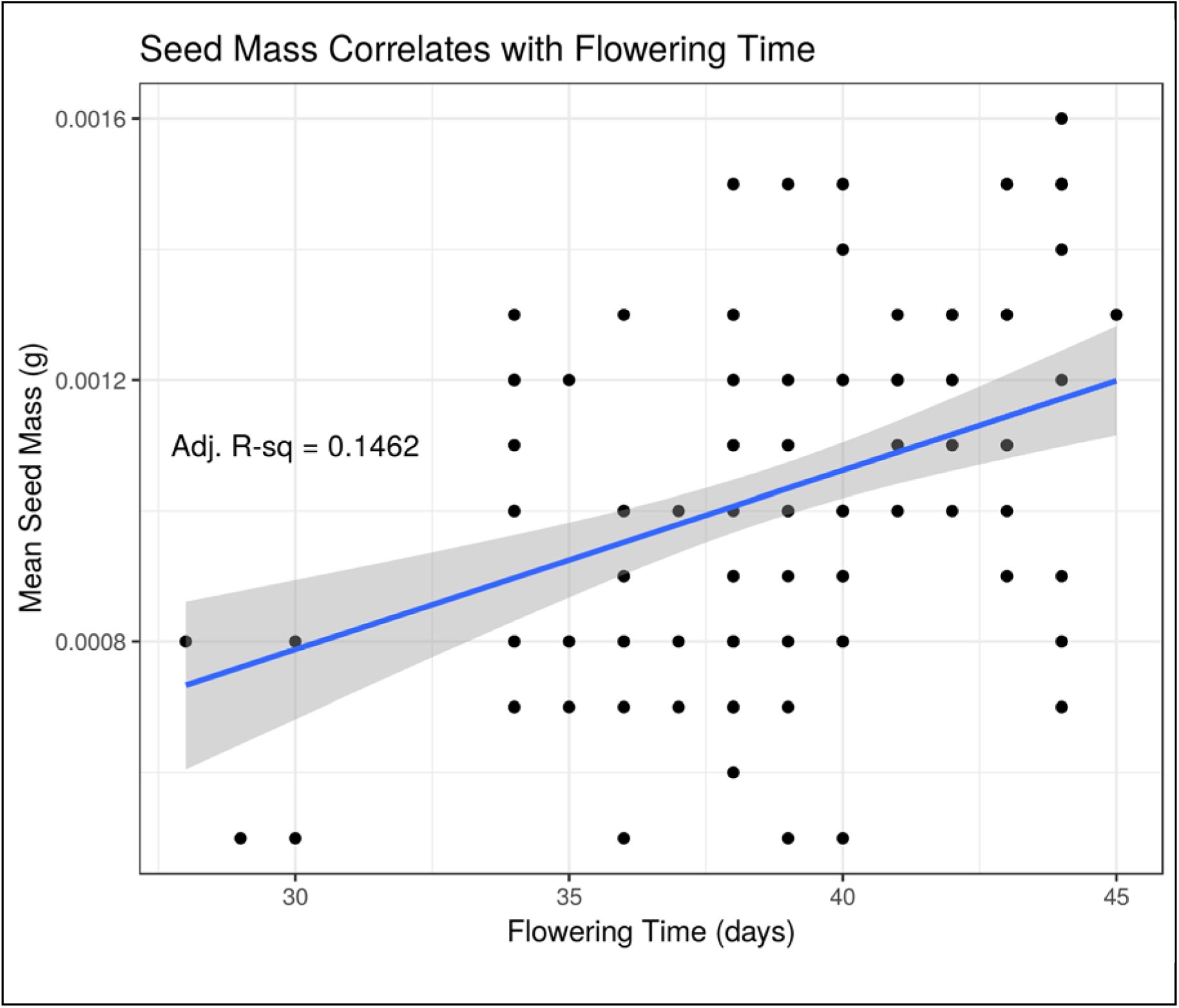

## LITERATURE CITED

Aleklett, K., M. Hart, and A. Shade. 2014. The microbial ecology of flowers: an emerging frontier in phyllosphere research. Botany 92: 253–266.

Arnold, A.E., L.C. MejÍA, D. Kyllo, E.I. Rojas, Z. Maynard, N. Robbins, and E.A. Herre. 2003. Fungal endophytes limit pathogen damage in a tropical tree. Proceedings of the National Academy of Sciences 100: 156491–15654.

Banta, J.A., I.M. Ehrenreich, S. Gerard, L. Chou, A. Wilczek, J. Schmitt, P.X. Kover, and M.D. Purugganan. 2012. Climate envelope modelling reveals intraspecific relationships among flowering phenology, niche breadth and potential range size in Arabidopsis thaliana. Ecology letters 15: 769–777.

Beattie, G.A. 2011. Water Relations in the Interaction of Foliar Bacterial Pathogens with Plants. Annual Review of Phytopathology 49: 533–555.

Berg, M., and B. Koskella. 2018. Nutrient-and Dose-Dependent Microbiome-Mediated Protection against a Plant Pathogen. Current Biology 28: 2487–2492.e3.

Bertrand, A., D. PrÉVost, F.J. Bigras, and Y. Castonguay. 2007. Elevated Atmospheric CO 2 and Strain of Rhizobium Alter Freezing Tolerance and Cold-induced Molecular Changes in Alfalfa (Medicago sativa). Annals of Botany 99: 275–284.

Busby, P.E., M. Ridout, and G. Newcombe. 2015. Fungal endophytes: modifiers of plant disease. Plant Molecular Biology 90: 645–655.

Cullings, K.W., and D.R. Vogler. 1998. A 5.8S nuclear ribosomal RNA gene sequence database: applications to ecology and evolution. Molecular Ecology 7: 919–923.

Davitt, A.J., M. Stansberry, and J.A. Rudgers. 2000. Do the costs and benefits of fungal endophyte symbiosis vary with light availability? New Phytologist 188: 824–834.

Doty, S.L. 2017. Functional Importance of the Plant Endophytic Microbiome: Implications for Agriculture, Forestry, and Bioenergy. In Functional Importance of the Plant Microbiome, 1–5. Springer, Cham. Available at: https://link.springer.com/chapter/10.1007/978-3-319-65897-1_1 [Accessed June 20, 2018].

Edwards, J., C. Johnson, C. Santos-MedellÍN, E. Lurie, N.K. Podishetty, S. Bhatnagar, J.A. Eisen, and V. Sundaresan. 2015. Structure, variation, and assembly of the root-associated microbiomes of rice. Proceedings of the National Academy of Sciences 112: E911–E920.

Egamberdieva, D., S.J. Wirth, A.A. Alqarawi, E.F. Abd_Allah, and A. Hashem. 2017. Phytohormones and Beneficial Microbes: Essential Components for Plants to Balance Stress and Fitness. Frontiers in Microbiology 8:. Available at: https://www.frontiersin.org/articles/10.3389/fmicb.2017.02104/full [Accessed June 20, 2018].

Falconi, C.J., and K. Mendgen. 1994. Epiphytic fungi on apple leaves and their value for control of the postharvest pathogens Botrytis cinerea, Monilinia fructigena and Penicillium expansum/Epiphytische Pilze auf Apfelblättern und ihre Eignung für die Bekämpfung der Apfelfäuleerreger Botrytis cinerea, Monilinia fructigena und Penicillium expansum. Zeitschrift für Pflanzenkrankheiten und Pflanzenschutz/Journal of Plant Diseases and Protection 38–47.

Fox, J., and S. Weisberg. 2011. An {R} Companion to Applied Regression. 2nd ed. Sage, Thousand Oaks, CA, USA. Available at: http://socserv.socsci.mcmaster.ca/jfox/Books/Companion.

Henning, J.A., D.J. Weston, D.A. Pelletier, C.M. Timm, S.S. Jawdy, and A.T. Classen. 2016. Root bacterial endophytes alter plant phenotype, but not physiology. PeerJ 4: e2606.

Lau, J.A., and J.T. Lennon. Evolutionary ecology of plant–microbe interactions: soil microbial structure alters selection on plant traits. New Phytologist 192: 215–224.

Lundberg, D.S., S.L. Lebeis, S.H. Paredes, S. Yourstone, J. Gehring, S. Malfatti, J. Tremblay, et al. 2012. Defining the core Arabidopsis thaliana root microbiome. Nature 488: 86–90.

Mendes, L.W., E.E. Kuramae, A.A. Navarrete, J.A. Van Veen, and S.M. Tsai. 2014. Taxonomical and functional microbial community selection in soybean rhizosphere. The ISME Journal 8: 1577–1587.

Panke-Buisse, K., A.C. Poole, J.K. Goodrich, R.E. Ley, and J. Kao-Kniffin. 2015. Selection on soil microbiomes reveals reproducible impacts on plant function. The ISME Journal 9: 980–989.

Peay, K.G., M. Belisle, and T. Fukami. 2011. Phylogenetic relatedness predicts priority effects in nectar yeast communities. Proceedings of the Royal Society of London B: Biological Sciencesrspb20111230.

Perrine-Walker, F.M., E. Gartner, C.H. Hocart, A. Becker, and B.G. Rolfe. 2007. Rhizobium-Initiated Rice Growth Inhibition Caused by Nitric Oxide Accumulation. Molecular Plant-Microbe Interactions 20: 283–292.

Pozo, M.I., M.-A. Lachance, and C.M. Herrera. 2012. Nectar yeasts of two southern Spanish plants: the roles of immigration and physiological traits in community assembly. FEMS Microbiology Ecology 80: 281–293.

Preston, G.M. 2017. Profiling the extended phenotype of plant pathogens. Molecular Plant Pathology 18: 443–456.

Ritpitakphong, U., L. Falquet, A. Vimoltust, A. Berger, J.-P. MÉTraux, and F. L’Haridon. 2016. The microbiome of the leaf surface of Arabidopsis protects against a fungal pathogen. New Phytologist 210: 1033–1043.

Rosas, U., Y. Mei, Q. Xie, J.A. Banta, R.W. Zhou, G. Seufferheld, S. Gerard, et al. 2014. Variation in Arabidopsis flowering time associated with cis-regulatory variation in CONSTANS. Nature Communications 5: 3651.

Rossmann, M., S.W. Sarango-Flores, J.B. Chiaramonte, M.C.P. Kmit, and R. Mendes. 2017. Plant Microbiome: Composition and Functions in Plant Compartments. In The Brazilian Microbiome, 7–20. Springer, Cham. Available at: https://link.springer.com/chapter/10.1007/978-3-319-59997-7_2 [Accessed June 19, 2018].

Saari, S., M. Helander, S.H. Faeth, and K. Saikkonen. 2010. The Effects of Endophytes on Seed Production and Seed Predation of Tall Fescue and Meadow Fescue. Microbial Ecology 60: 928–934.

Schlaeppi, K., and D. Bulgarelli. 2014. The Plant Microbiome at Work. Molecular Plant-Microbe Interactions 28: 212–217.

Vandenkoornhuyse, P., A. Quaiser, M. Duhamel, A. Le Van, and A. Dufresne. 2015. The importance of the microbiome of the plant holobiont. New Phytologist 206: 1196–1206.

Wagner, M.R., D.S. Lundberg, D. Coleman-Derr, S.G. Tringe, J.L. Dangl, and T. Mitchell-Olds. 2014. Natural soil microbes alter flowering phenology and the intensity of selection on flowering time in a wild Arabidopsis relative. Ecology letters 17: 717–726.

Wagner, M.R., D.S. Lundberg, T.G. Del Rio, S.G. Tringe, J.L. Dangl, and T. Mitchell-Olds. 2016. Host genotype and age shape the leaf and root microbiomes of a wild perennial plant. Nature Communications 7: 12151.

Westoby, M., E. Jurado, and M. Leishman. 1992. Comparative evolutionary ecology of seed size. Trends in Ecology & Evolution 7: 368–372.

Wickham, H. 2009. ggplot2: Elegant Graphics for Data Analysis.Springer-Verlag, New York.

Wolfe, B.E. 2018. Using Cultivated Microbial Communities To Dissect Microbiome Assembly: Challenges, Limitations, and the Path Ahead. mSystems 3: e00161–17.

Zahn, G., and A.S. Amend. 2017. Foliar microbiome transplants confer disease resistance in a critically-endangered plant. PeerJ 5: e4020.

