## Supplementary material for "Foliar fungi alter reproductive timing and allocation in *Arabidopsis* under normal and water-stressed conditions": Raw Data and analysis code: Analysis_and_Figures.R

```
library(dplyr)
library(ggplot2)
library(vegan)
library(multcompView)

'%ni%' = Negate('%in%')


#### read in and prep data ####
dat = read.csv(file = "Spreadsheets/Arabidopsis_Data_Sheet_for_R.csv", header = TRUE)
dat$Germination_Date = as.POSIXct(dat$Germination_Date, format = '%m-%d-%Y')
dat$Inoculation_Date = as.POSIXct(dat$Inoculation_Date, format = '%m-%d-%Y')
dat$Flowering_Date = as.POSIXct(dat$Flowering_Date, format = '%m-%d-%Y')


#add flowering times (days) to dat
dat = dat %>%
  mutate(Days_to_Flower = Flowering_Date - Germination_Date)

dat$Days_to_Flower = as.numeric(dat$Days_to_Flower)


# factor Inoculation richness
dat$InocRichness = factor(dat$InocRichness)

#remove army samples
dat = dat[1:160,]
dat$Fungal_Treatment = factor(dat$Fungal_Treatment)

# add color for plot
dat$Group = NA
dat$Group[dat$InocRichness !="0"] <- "Inoculated"
dat$Group[dat$InocRichness =="0"] <- "Sterile"


#### Plot of richness vs flower time ####
ggplot(dat, aes(x=InocRichness,y=Days_to_Flower, fill=Group)) +
geom_boxplot() + theme_bw() +
  labs(y="Days to Flower",x="Inoculum Richness") +
  ggtitle("Sterile plants had consistently shorter flowering times")
ggsave("Figures/SI_Fig_1.tiff", dpi = 300)


#### Some Stats ####

###     DO THIS WITH ACTUAL COMMUNITY RESULTS INSTEAD OF JUST "SPRAYED TREATMENT" AND IT WILL BE MORE MEANINGFUL!!!! ###
#anova on flowering times
aov_flower_1 = aov(data = dat[which(dat$Water == "Norm"),], Days_to_Flower ~ Fungal_Treatment)
aov_flower_2 = aov(data = dat[which(dat$Water == "Low"),], Days_to_Flower ~ Fungal_Treatment)

#post-hoc tukey test
tukey_flower_1 = TukeyHSD(aov_flower_1)
tukey_flower_2 = TukeyHSD(aov_flower_2)

#extract p-values from tukeyhsd output ($Fungal_Treatment[,4]) and move into a data frame
tukey_flower_1_pvals = as.data.frame(tukey_flower_1$Fungal_Treatment[,2-4])
tukey_flower_2_pvals = as.data.frame(tukey_flower_2$Fungal_Treatment[,2-4])

#add pairings to df
tukey_flower_1_pvals$Pair = row.names(tukey_flower_1_pvals)
tukey_flower_2_pvals$Pair = row.names(tukey_flower_2_pvals)


#rename and sort
names(tukey_flower_1_pvals) <- c("Diff","Upper","P_val","Pair")
Flower_time_sig.diff.Norm = tukey_flower_1_pvals %>%
  arrange(Diff) %>%
  filter(P_val < 0.05) %>%
  select(Pair, Diff, P_val)

names(tukey_flower_2_pvals) <- c("Diff","Upper","P_val","Pair")
Flower_time_sig.diff.Low = tukey_flower_2_pvals %>%
  arrange(Diff) %>%
  filter(P_val < 0.05) %>%
  select(Pair, Diff, P_val)

write.csv(Flower_time_sig.diff.Norm, file = "Spreadsheets/Arabid_Flowering_Time_Sig_Diffs_Norm.csv",
          row.names = FALSE)
write.csv(Flower_time_sig.diff.Low, file = "Spreadsheets/Arabid_Flowering_Time_Sig_Diffs_Low.csv",
          row.names = FALSE)


####  Community presence matrix ####

A = dat[dat$A==1,]
B = dat[dat$B==1,]
C = dat[dat$C==1,]
D = dat[dat$D==1,]
N = dat[dat$NEG==1,]


# Plot fungal isolate presence vs days to flower ####
A.fig = ggplot() +
  geom_violin(aes(x="A",y=A$Days_to_Flower), fill="Gray1") +
  geom_violin(aes(x="B",y=B$Days_to_Flower), fill="Gray1") +
  geom_violin(aes(x="C",y=C$Days_to_Flower), fill="Gray1") +
  geom_violin(aes(x="D",y=D$Days_to_Flower), fill="Gray1") +
  geom_violin(aes(x="N",y=N$Days_to_Flower), fill="Gray") +
  lims(y=c(0,50)) +
  labs(x="Fungi included in treatment", y= "Days to Flower") +
  ggtitle("Fungal isolate identity was less important than presence of any fungi") +
  theme_bw()
A.fig
ggsave("Figures/Fig_2.tiff", dpi=300, height = 8, width = 10)


# permanova with spp matrix as response ####
veg.matrix = (dat[,c("A","B","C","D","NEG")])
adonis(veg.matrix ~ dat$Days_to_Flower+dat$Water)

# without negative controls
no.neg.veg.matrix = (dat[dat$Fungal_Treatment != "NEG",c("A","B","C","D")])
no.neg.dat = dat[dat$Fungal_Treatment != "NEG",]
adonis(no.neg.veg.matrix ~ no.neg.dat$Days_to_Flower+no.neg.dat$Water)

#### Competition Results ####

# read ImageJ data
comp = read.delim("Spreadsheets/COMPETITION_RESULTS.csv", sep = "\t")[,1:4]

# remove empty rows
comp = na.omit(comp)

# convert pixels to cm2 with known scale (8462322 pixels in 5cm dish at ImageJ measurement scale)
comp$Area.cm2 = (comp$Area.Pxl * 19.63) / 8462322

# set baselines for each measurement day
for(i in c("A","B","C","D")){
  assign(paste("mean.4.",i,sep = ""), (mean(comp[comp$Treatment == i & comp$Day == 4,'Area.cm2'])), envir = .GlobalEnv)
}

for(i in c("A","B","C","D")){
  assign(paste("mean.9.",i,sep = ""), (mean(comp[comp$Treatment == i & comp$Day == 9,'Area.cm2'])), envir = .GlobalEnv)
}

# get baseline growth rates for pure cultures (cm2 per day)
Rate_A = (mean.9.A - mean.4.A) / 5
Rate_B = (mean.9.B - mean.4.B) / 5
Rate_C = (mean.9.C - mean.4.C) / 5
Rate_D = (mean.9.D - mean.4.D) / 5

# compare growth rates of fungi in competition to baseline rates
Comp.A = comp[comp$Component == 'A',] 
Comp.B = comp[comp$Component == 'B',] 
Comp.C = comp[comp$Component == 'C',] 
Comp.D = comp[comp$Component == 'D',] 

Comp.A.4 = comp[comp$Component == 'A' & comp$Day == 4,] 
Comp.A.9 = comp[comp$Component == 'A' & comp$Day == 9,]
Comp.B.4 = comp[comp$Component == 'B' & comp$Day == 4,] 
Comp.B.9 = comp[comp$Component == 'B' & comp$Day == 9,]
Comp.C.4 = comp[comp$Component == 'C' & comp$Day == 4,] 
Comp.C.9 = comp[comp$Component == 'C' & comp$Day == 9,]
Comp.D.4 = comp[comp$Component == 'D' & comp$Day == 4,] 
Comp.D.9 = comp[comp$Component == 'D' & comp$Day == 9,]

Comp.A$Rate = (Comp.A.9$Area.cm2 - Comp.A.4$Area.cm2) / 5
Comp.B$Rate = (Comp.B.9$Area.cm2 - Comp.B.4$Area.cm2) / 5
Comp.C$Rate = (Comp.C.9$Area.cm2 - Comp.C.4$Area.cm2) / 5
Comp.D$Rate = (Comp.D.9$Area.cm2 - Comp.D.4$Area.cm2) / 5

Comp.A$Relative_Rate = Comp.A$Rate / Rate_A
Comp.B$Relative_Rate = Comp.B$Rate / Rate_B
Comp.C$Relative_Rate = Comp.C$Rate / Rate_C
Comp.D$Relative_Rate = Comp.D$Rate / Rate_D

a = Comp.A %>% group_by(Treatment) %>% summarise(mean.relative.growth = mean(Relative_Rate))
b = Comp.B %>% group_by(Treatment) %>% summarise(mean.relative.growth = mean(Relative_Rate))
c = Comp.C %>% group_by(Treatment) %>% summarise(mean.relative.growth = mean(Relative_Rate))
d = Comp.D %>% group_by(Treatment) %>% summarise(mean.relative.growth = mean(Relative_Rate))

a$Isolate = "A"
b$Isolate = "B"
c$Isolate = "C"
d$Isolate = "D"

Rel.Growth = as.data.frame(rbind(a,b,c,d))
Full.Rel.Growth = as.data.frame(rbind(Comp.A,Comp.B,Comp.C,Comp.D))
names(Full.Rel.Growth)[3] <- "Isolate"


# plot culture competitions ####

# removed a,b,c,d pure cultures
ggplot(Full.Rel.Growth, mapping = aes(x = Treatment, y = Relative_Rate, col = Isolate)) +
  geom_hline(yintercept = 1, linetype = 2) +
  geom_boxplot() + theme_bw() + ggtitle("Relative Growth Rates of Isolates in competition") +
  labs(subtitle = "Points above 1 indicate faster growth in a respective competition than in pure culture") +
  geom_vline(xintercept = c(1.5,2.5,3.5,4.5,5.5,6.5,7.5,8.5,
                            9.5,10.5), colour = "Gray") +
  theme(panel.grid.major.x = element_blank()) +
  scale_x_discrete(limits = c("AB","AC","AD","BC","BD","CD","ABC","ABD","ACD",
                              "BCD","ABCD"))
ggsave("Figures/Fig_3.tiff", dpi = 300, height = 8, width = 10)

# look at pairs (means in paired competitions) ####
pairs = c("AB","AC","AD","BC","BD","CD")

# write paired growth data to results file 

sink("Tables/Paired_Competition_Means.txt")
for(i in pairs[]){
  Obs =  comp %>%
    filter(Treatment == i & Day == 9) %>%
    group_by(Component) %>%
    summarise(N = n(), Mean = mean(Area.Pxl), Treatment = i)
  print(Obs)
}
sink(NULL)


# Flowering time Plot ####

library(plyr)
# Define interaction levels by group combination
dat.Int <- ddply(dat,.(Fungal_Treatment,Water),summarise, val = mean(Days_to_Flower))

ggplot(dat, aes(x=Fungal_Treatment,y=Days_to_Flower,fill=Water)) +
  geom_boxplot() + 
  geom_line(data = dat.Int, aes(y = val, group = Water, col = Water)) + # add interaction line
  theme_bw()

# Find difference between norm and low water flowering times for each fungal treatment:

stacked = dat.Int
stacked$Fungal_Treatment = as.character(stacked$Fungal_Treatment)
stacked$Water = as.character(stacked$Water)
diff.water = as.data.frame(t(unstack(stacked, val ~ Fungal_Treatment)))
diff.water$diff = diff.water$V2 - diff.water$V1
# order to match other plots
diff.water = diff.water[c("A","B","C","D","AB","AC","AD","BC","BD","CD","ABC","ABD","ACD",
                          "BCD","ABCD","NEG"),]

mean.norm = mean(dat[dat$Water == "Norm","Days_to_Flower"])
mean.low = mean(dat[dat$Water == "Low","Days_to_Flower"])

# annotate with difference in mean flowering time between low and normal water conditions
ggplot(dat, aes(x=Fungal_Treatment,y=Days_to_Flower,fill=Water)) +
  geom_hline(yintercept = mean.norm, colour = "Cyan4", linetype = 2) +
  geom_hline(yintercept = mean.low, colour = "IndianRed3", linetype = 2) +
  geom_boxplot() + 
  geom_line(data = dat.Int, aes(y = val, group = Water, col = Water)) + # add interaction line
  theme_bw() +
  annotate("text", x=c(1:nlevels(dat$Fungal_Treatment)),
           y=26,label=signif(diff.water$diff,2),col = "DarkGreen") + 
  ggtitle("Flowering Time (days) by Epiphyte and Water Treatment") +
  labs(x = "Fungal Treatment", y= "Days to Flower") +
  scale_x_discrete(limits = c("A","B","C","D","AB","AC","AD","BC","BD","CD","ABC","ABD","ACD",
                              "BCD","ABCD","NEG")) +
  geom_vline(xintercept = c(1.5,2.5,3.5,4.5,5.5,6.5,7.5,8.5,
                            9.5,10.5,11.5,12.5,13.5,14.5,15.5), colour = "Gray") +
  theme(panel.grid.major.x = element_blank()) +
  annotate("text", x=8.5,y=27.5,label="Differences between mean flowering times (days) under altered watering regimes:", col = "DarkGreen", cex = 6) # this is sort of a proxy for how much stress the plant was under due to water decrease

ggsave("~/Desktop/Arabidopsis/R_PROJECT/Figures/Fig_1.tiff", dpi = 300, height = 10, width = 12)    

# Plot seed mass as function of flowering time ####
# correlate seed mass to flower time
summary(lm(dat$Seed_Mass_g ~ dat$Days_to_Flower))

Fig_SI2 = ggplot(mapping = aes(x=dat$Days_to_Flower,y=(dat$Seed_Mass_g))) +
  geom_point() +
  geom_smooth(method = "lm") +
  theme_bw() + labs(x="Flowering Time (days)",y="Mean Seed Mass (g)") + ggtitle("Seed Mass Correlates with Flowering Time") +
  annotate("text",x=30,y=0.0011,label = "Adj. R-sq = 0.1462")
ggsave("./Figures/SI_Fig_2.tiff", dpi = 300)
```
